# Attention modulates EEG spectral activity across serial positions

**DOI:** 10.64898/2026.01.01.697233

**Authors:** Jacqueline S. Kim, Adam W. Broitman, Khena M. Swallow

**Author notes:** To whom correspondence should be addressed: Adam Wood Broitman University of Pennsylvania, Dept. of Psychology 425 S University Ave, Levin 205 Philadelphia, PA, 19104.

## Abstract

Electroencephalographic (EEG) activity has been associated with attentional processes such as task-irrelevant stimulus suppression and orienting. The aim of this study was to understand how spectral activity evolves as words are presented in a word list learning task and as multiple components of attention are engaged under multitasking demands. We recorded scalp EEG in a delayed free-recall task in which participants encoded common noun word lists either under full attention (encoding only) or divided attention (simultaneous target detection task). In the target detection task, participants pressed a response button when a word was paired with a target-colored square but not for words paired with a distractor-colored square (cue condition). Activity in the high gamma frequencies (50-100 Hz) decreased while alpha activity (8-12 Hz) increased throughout word list presentation (i.e., across serial positions), consistent with prior work demonstrating effects of cognitive load on EEG spectral power. Divided attention attenuated this change in gamma power across the list. Cue condition modulated alpha activity evenly across serial positions, with consistently lower power during target trials than distractor trials. Our findings suggest that frequency-specific characteristics of processing during list encoding may reflect multiple components of attention that vary over different time scales. Gamma frequencies capture the interactive effects of cognitive load and dual-task demands, while alpha power separately reflects gradual increases in cognitive load and momentary orienting.

## Introduction

Attention to external information fluctuates in response to daily life’s demands, affecting how information is processed and represented in the brain (Chun & Turk-Browne, 2007; Kuo et al., 2012; Robinson, 1995). Cognitive load and task demands influence information processing over multiple timescales, and they may be captured in spectral components of scalp electroencephalographic (EEG) activity. In particular, greater spectral power in low frequency bands (e.g., alpha, 8-12 Hz) often corresponds to inhibited processing (Pereda et al., 1998; Roux et al., 2022). Power in high frequency bands (e.g., high gamma, 50-100 Hz) has been linked to increased cognitive processing (Miller et al., 2018). However, the degree to which alpha and high gamma band power reflect different components of attention and their interactions remains unclear. To better characterize how various aspects of attention influence neural processing over time, we examined the interactive effects of cognitive load with divided attention and orienting on spectral components of scalp EEG during a word list learning task.

Neural activity reflects changes in multiple components of attention during task performance. One such component is cognitive load (Kahneman, 1973; Lavie, 2005; Posner & Boies, 1971), broadly defined as the amount of information the brain must retain or process at a given moment (Cabañero Gómez et al., 2021). It increases with the number of items maintained in memory (Cabañero Gómez et al., 2021; Doherty & Logie, 2016) and the complexity of the task performed (i.e., number of steps or processes that must be coordinated; Zeitlhofer et al., 2024). Gamma frequency band power is modulated by cognitive load. It increases with task-relevant attention (Fries et al., 2001), working memory load (Howard et al., 2003; Van Vugt et al., 2010), task complexity (Simos et al., 2002), and cortical activation (Engel et al., 2001; Gruber et al., 1999; Merker, 2013; Ray & Cole, 1985; Tiitinen et al., 1993). Alpha band power is also associated with cognitive load, though both increases and decreases may be observed (Jensen et al., 2002; Randazzo et al., 2019). This may reflect its potential role in inhibiting processing (and gamma activity) that interferes with task performance (Klimesch, 2012). For example, alpha band power decreases in regions that represent items held in working memory, while increasing elsewhere (Jokisch & Jensen, 2007). Gamma and alpha activity may also capture momentary changes in task demands. Alpha band power reliably decreases shortly after the appearance of target stimuli in detection and visual search tasks, consistent with orienting attention toward behaviorally relevant stimuli (de Graaf et al., 2020; He et al., 2021; Klimesch et al., 1998). Similarly, gamma power can increase during brief periods of active processing, reflecting localized cortical activation (DeHaan et al., n.d.; Fries, 2009). Together, this work suggests that sustained increases in cognitive load (e.g., due to multitasking) and momentary shifts in task demands (e.g., due to target detection) may jointly impact spectral EEG activity at multiple timescales. In this case, we may be able to observe differential effects of cognitive load and orienting on spectral EEG activity.

This study investigates the effects of sustained and transient aspects of attention on gamma and alpha band activity during word list learning. Both alpha and gamma band activity change over serial positions as a list of words is presented. Alpha-band activity increases over the course of the list while gamma band activity decreases (Sederberg et al., 2006; Serruya et al., 2014). The effect of word list serial position on gamma activity has been termed the *gamma primacy effect*. Changes in gamma and alpha power with serial position are hypothesized to reflect reductions in the ability to process new words due to increased cognitive load or divided attention between list items (Sederberg et al., 2006). Consistent with this possibility, the ability to encode newly presented words may decrease with serial position and greater cognitive load (Azizian & Polich, 2007; Portrat et al., 2008), potentially leading to primacy effects in encoding (Bennet, 1962; Page & Norris, 1998; Serruya et al., 2014; Tulving, 2007).

We examined spectral power in list learning under full attention and in a continuous dual-task encoding paradigm commonly used to study the attentional boost effect (ABE). Divided attention and attentional orienting were introduced by pairing an encoding task with momentary, salient cues requiring a response (targets) or cues that were to be ignored (distractors). In the attentional boost effect, items presented with targets are better recalled than items presented with distractors (Broitman & Swallow, 2024; Mulligan et al., 2014; Swallow & Jiang, 2010). Targets also evoke stronger reductions in alpha power, particularly when the coinciding item is recalled (Broitman & Swallow, 2025).

While prior studies have investigated gamma and alpha power during dual-tasks (Broitman et al., 2025; Broitman & Swallow, 2025; Long & Kahana, 2017), none have tracked how this spectral activity may evolve with multitasking demands across serial positions. Given the Sederberg et al. (2006) proposal that the gamma primacy effect is due to the transition from focused to divided encoding states with increasing list items, we expected that introducing a second task would suppress it. Increases in alpha power with serial position may also reflect increased cognitive load and divided attention. If so, alpha may also show interactive effects of dual-tasking with serial position. However, changes in alpha power over serial positions could reflect reduced attention to the words, possibly limiting any interactive effects of dual-tasking and serial position on neural activity.

In addition to introducing dual-task demands, the detection task also required participants to respond to some cues (targets) but not others (distractors), creating transient changes in task demands. Gamma power in list learning may reflect sustained, rather than transient aspects of attention (Broitman & Swallow, 2025; Engel et al., 2001; Gruber et al., 1999) and thus may be relatively stable across cue conditions. In contrast, alpha power reliably decreases following target detection (de Graaf et al., 2020; Klimesch et al., 1998) and therefore should be lower on target trials. However, whether this effect interacts with those of cognitive load is unknown.

## Methods

### Participants

The data from this study were previously analyzed and reported in Broitman & Swallow (2025), where further detail can be found. The research protocol was approved by the Cornell University Institutional Review Board. Young adults (aged 18-40 years) were recruited around the Cornell University campus and the greater Ithaca area. They were given a choice of either $10 per hour or psychology course credit for compensation. All participants signed a physical informed consent document and were fully debriefed following the study.

Eligible participants were neurologically healthy, native English speakers with normal or corrected-to-normal vision, including normal color vision verified by the Hardy, Rand, and Rittler pseudoisochromatic color blindness test (citation). Four participants were excluded for not meeting performance criteria on the detection task (>80% hits, <10% false alarms) or recall task (>100, or 22%, words correctly recalled), resulting in a final sample size of 61 participants, 21 in the single task and 40 in the dual-task condition.

### Equipment and Materials

Participants were fitted with a 64-channel Biosemi ActiveTwo geodesic sensor cap, and EEG data was collected at a sampling rate of 512 Hz. Situated around 90 cm in front of a 17′′ CRT monitor (75Hz refresh rate) and keyboard, participants were asked to minimize blinking and body movements during recording. Additional details regarding word list construction and participant recruitment are provided in Broitman & Swallow (2025).

### Experiment

Each word list began with a fixation cross for 2000 ms. Each trial consisted of a noun displayed with two colored squares (red [RGB: 255, 0, 0] or green [RGB: 0, 255, 0]) above and below it for 100 ms. The word remained alone without the squares for an additional 400 ms before a variable interstimulus interval of 750-1250 ms. There were equal numbers of red and green-paired words within each list, evenly distributed across serial positions.

This cycle continued until every word within a list was presented. A math distractor task of simple arithmetic problems (e.g., 3 + 5 − 2 = ?) accommodated for the advantage to end-list items and followed encoding for 30 seconds. Participants were then asked to type as many words as they could recall, in any order, for 75 seconds.

#### Dual Task Conditions

40 participants were assigned to the *divided attention* condition. Each participant was presented with 28 lists, each list composed of 16 words, for a total of 448 words per participant. During the list presentation, participants were simultaneously tasked with pressing the spacebar for trials paired with target-colored squares only. Half of these participants were assigned red as their target color, while the other half responded to green targets. Subsequent analyses grouped serial positions into 4 groups: *early* (1-4), *early-middle* (5-8), *late-middle* (9-12), and *late* (13-16).

#### Single Task Conditions

21 participants were assigned to the *full attention* condition. Each participant was presented with 24 lists, each composed of 18 words, for a total of 432 words. Serial positions were again grouped into 4 groups: *early* (1-4), *early-middle* (5-9), *late-middle* (10-14), and *late* (15-18). In this full attention or single task condition, participants were told to ignore the colored squares presented with the words during the encoding period.

### EEG Data Preprocessing

The MNE software package (Gramfort et al., 2013) was used to preprocess the collected EEG recordings in multiple stages with Python scripts. A bandpass filter attenuated frequencies outside of 0.1-100 Hz, and a notch filter helped to eliminate environmental or powerline noise at 60 Hz. We processed the data using independent component analysis and then manually identified and removed components that were consistent with artifacts (e.g, from eyeblinks, heartbeats, muscle movements) based on visual inspection of scalp topography and temporal alignment with stereotyped artifacts. Finally, we used the package’s spherical spline interpolation method to replace channels displaying erratic activity or excessive electrical noise based on interpolation from nearby channels.

### Spectral Density Analyses

We performed time-frequency decomposition using a Morlet wavelet transformation with a wave number of 5. Power spectral density was calculated at 18 logarithmically spaced frequencies from 2 to 100 Hz. Power values were downsampled from the original rate of 512 Hz to 18 Hz and computed from -500 to 1000 ms relative to the onset of the word. We also applied a 1200 ms buffer on each side to account for edge effects. Powers were then Z-transformed within each channel and within each frequency, for each trial and participant, to normalize the data. Z-transformation entailed subtraction of the mean and division by the standard deviation for all power values for each trial, electrode, and frequency. Since power was Z-scored within each participant, there was no main effect of task condition (between participants). As we limited our analyses to post-trial times, data was averaged within the 0-1000 ms time window.

All analyses were restricted to the medial parietal region of interest (ROI) where alpha and gamma power correlates of attention were previously observed (Broitman & Swallow, 2025; Sederberg et al., 2006). This ROI included 13 electrodes: Pz, P1, P2, P3, P4, CPz, CP1, CP2, CP3, CP4, POz, PO1, and PO4.

Before further analyzing the power spectral densities (PSDs), powers were averaged within alpha (8-12 Hz) and high gamma (50-100 Hz) frequencies. Outlier trials with PSD values of |z| > 3 were removed.

### Statistical Analyses

Linear mixed effect models (LME) of each dependent variable (recall rates, alpha PSD, gamma PSD) were fit in R (R core team, 2021) using the lmer package (Bates et al., 2014). Participant and list number were included as random effects in all models. Serial position bin was modeled as a fixed effect with linear and quadratic terms. Separate models with additional fixed effects evaluated the effects of divided attention and cue condition. The effects of divided attention were evaluated by including task condition and its interaction with serial position bin as fixed effects. The effects of cue condition were evaluated in a model that was limited to dual-task participants and included cue condition and its interaction with serial position bin as fixed effects. These models took the general form of:

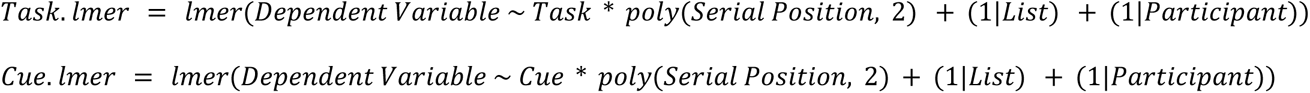

We characterized each model using ANOVA with the lmerTest package (Kuznetsova et al., 2017a). The emmeans package (Lenth et al., 2025) was used for contrast analyses and Holm corrections for multiple comparisons. Emtrends compared the change in gamma or alpha power across serial positions between either task condition (single or dual) or cue condition (target or distractor).

## Results

### Behavioral Results

Target-paired words were better recalled overall than distractor-paired words, replicating the attentional boost effect. Error bars represent standard errors. As reported in Broitman & Swallow (2025), dual-task participants accurately performed the detection task by quickly responding to targets and not distractors.

Recall rates were similar between task conditions (Figure 2a; *p* = .57). Consistent with prior analyses, a main effect of serial position bin indicated elevated recall rates at early and late list positions (primacy and recency effects) *F*(2, 25880) = 76, *p* <.001, η²_p_ = .01. However, an interaction of serial position bin and task condition indicated that recall remained more stable across list positions among dual-task participants than single-task participants, F(23, 25880) = 8.0, *p* < .001, η²_p_ < .001. Post-hoc contrast analyses indicated that this interaction primarily reflected stronger recency effects among single-task participants, z = -2.43, *p* = 0.02, though these participants also demonstrated numerically greater primacy effects, z = -.4, *p* = 0.70. Among dual-task participants, we observed main effects of cue condition (target > distractor), *F*(1, 16876) = 78, *p* <.001, η²_p_ = .01, and serial position bin (primacy and recency) *F*(3, 16872) = 41, *p* <.001, η²_p_ = .01, but their interaction was not significant, *p* = .23 (Figure 2b). Thus, the additive effect of target detection on recall did not significantly vary across serial position bins.

**Figure 1.**
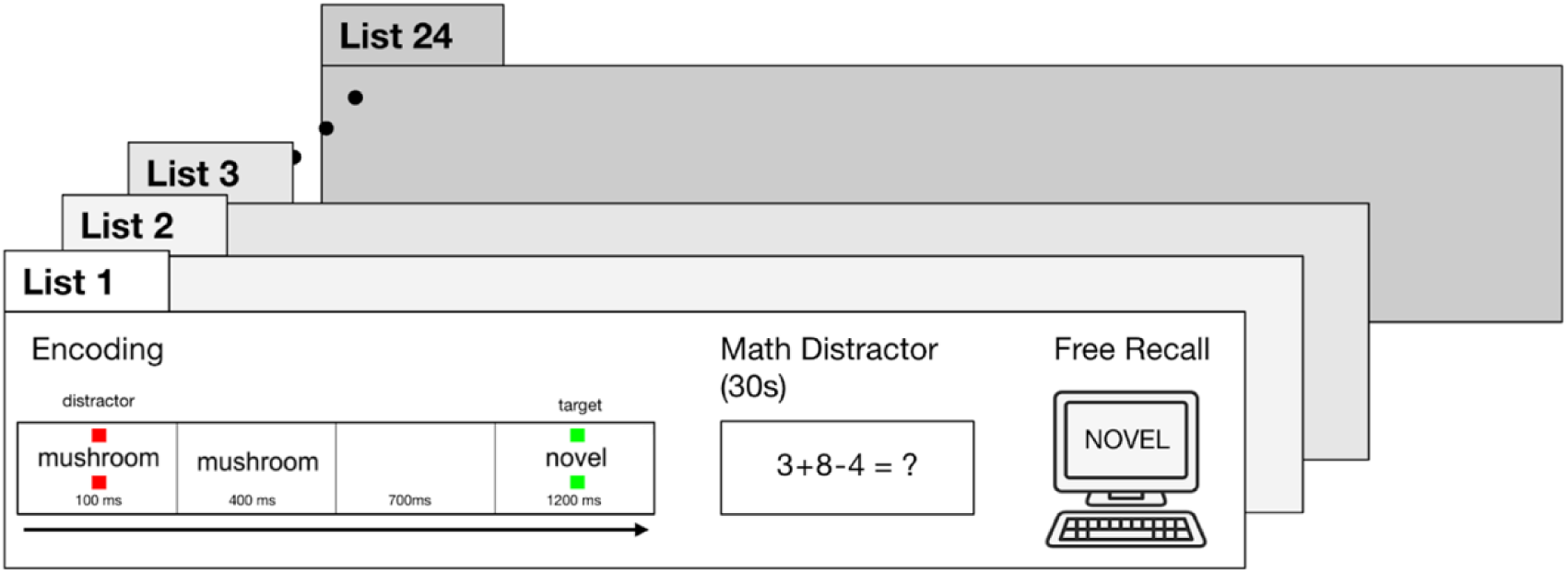
Experimental Task. Participants performed a delayed free recall task in cycles consisting of wordlist encoding, a 30-second math distractor task, and free recall (Figure 1). In the single-task condition, there were 24 lists (cycles) of 18 words each. In the dual-task condition, there were 28 lists of 16 words each.

**Figure 2.**
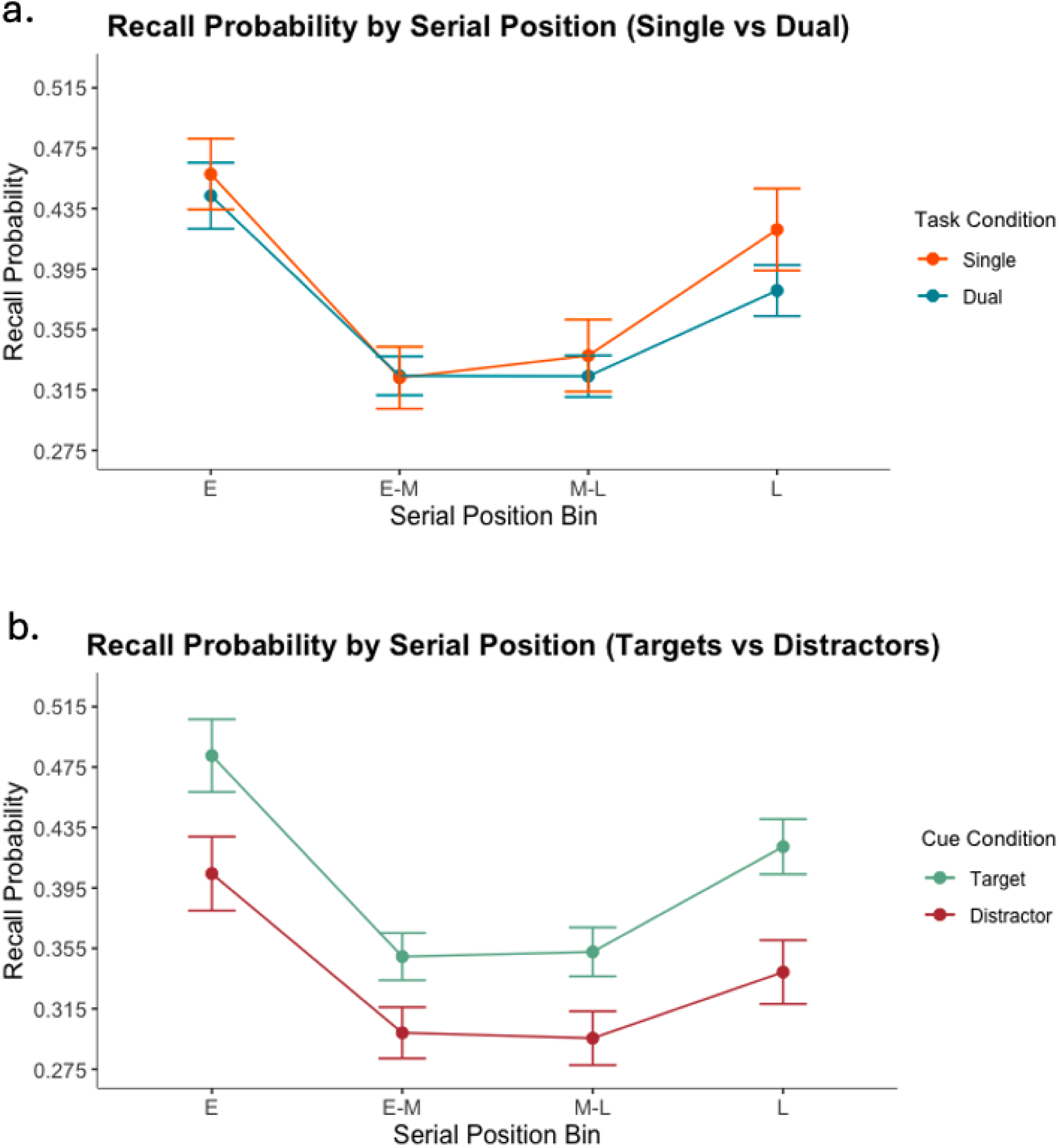
Serial Position Curve Between Task Conditions and Trial Types. Both graphs show the serial position effect, with a greater probability of recall at early serial positions (‘primacy effect’) and end-list positions (‘recency effect’). Panel (a) compares single and dual-task conditions, while (b) compares target and distractor cue conditions within dual-task trials.

### EEG Results

Because we reported main effects of task and cue condition in prior work (Broitman & Swallow, 2025), we report only main effects of and interactions with serial position bin.

#### Gamma Power Across Task Conditions

As shown in Figure 3a, we replicated the gamma primacy effect with increased gamma power at early list positions, as indicated by a main effect of serial position bin in our high gamma model, *F*(2, 25,507) = 121.67, *p* <.001, η²_p_ = .01. Further contrast analyses indicated gamma power was highest at the early serial position bin compared to all other bins, all z’s > 12.326, p’s < .001. Gamma was also higher in the early-middle bin compared to middle-late and late, z > 3.33, *p* < .05, though the middle-late bin did not significantly differ from the late bin, z = -1.33, *p* > 0.183. Gamma power therefore plateaued about mid-way through the lists.

**Figure 3.**
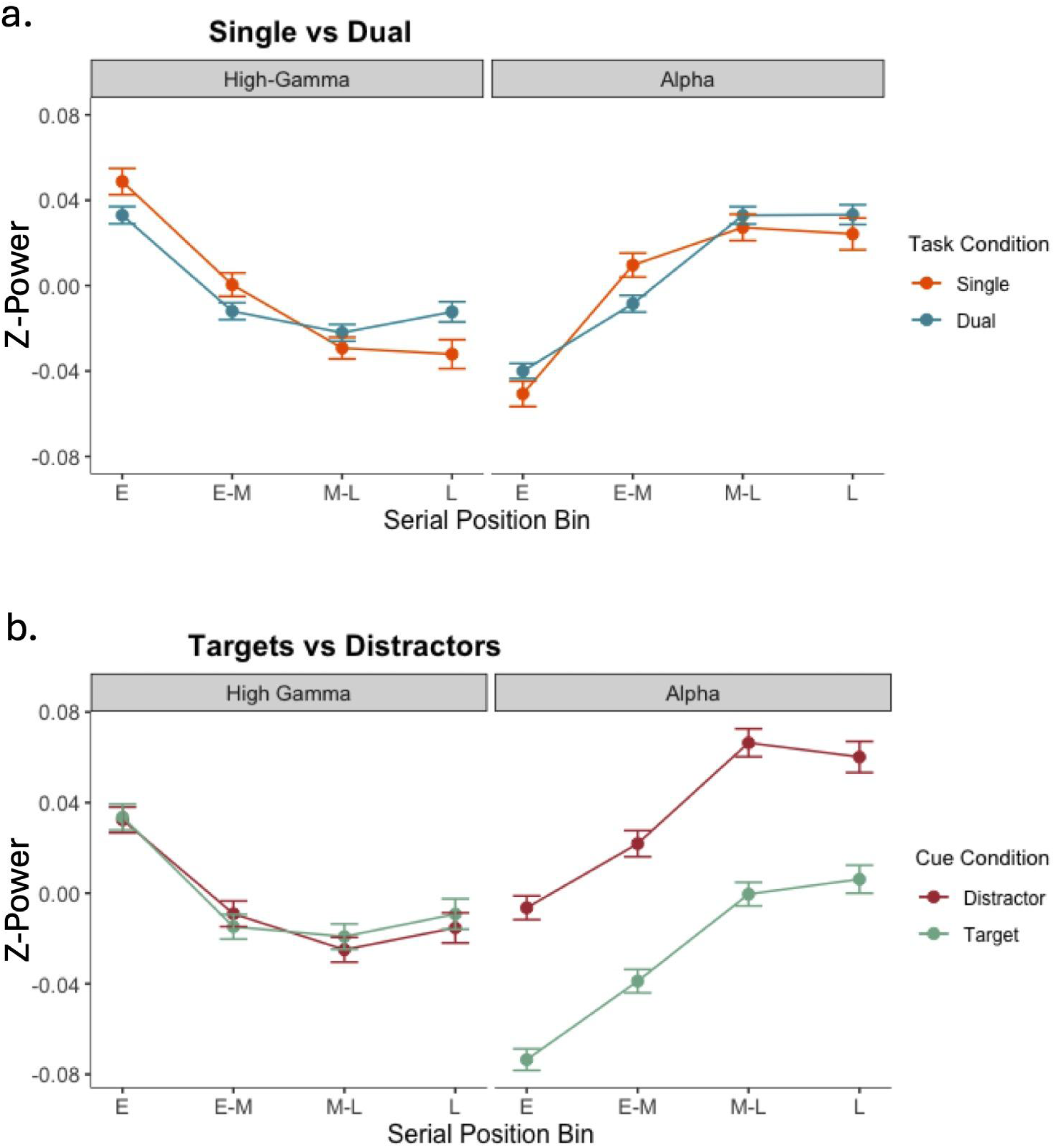
Comparing Task Condition and Cue Condition Effects Across Serial Positions on Alpha and High Gamma Power. We examined how alpha and high gamma spectral power varied with cognitive load during word-list encoding in an ABE paradigm. Panel (a) compares the effect of task condition (single or dual-task) across serial position. While the reductions in gamma over serial position bins (GPE) was smaller for dual-task trials, there was no effect of task condition on alpha power. (b) Effect of cue condition (target or distractor) across serial position. While there was no effect of cue condition across serial positions for high gamma, distractor trials displayed larger alpha power across list conditions for alpha. Error bars represent 95% confidence intervals.

The gamma primacy effect differed across task conditions, resulting in a significant interaction between serial position bin and task condition, *F*(2, 25,507) = 7.38, *p* <.001, η²_p_ = .0004. As depicted in Figure 3a, gamma power was initially higher and declined more rapidly in the full attention condition while multitasking showed lesser reduction over serial positions.We characterized the change in gamma power across serial positions using *emtrends*. Gamma power decreased more steeply in the single-task condition, *slope* = −0.031, *SE* = 0.003, compared to the dual-task condition, *slope* = −0.02, *SE* = 0.002. This difference in slopes was statistically significant, *z* = −3.54, *p* <.001. Additional contrast analyses were consistent with these results.

#### Gamma Across Cue Conditions

To explore whether the GPE differed by cue condition (target or distractor), we constructed an additional model restricted to dual-task trials. We observed a significant main effect of serial position bin, *F*(2, 16763) = 66.91, *p* <.001, η²_p_ = .007, confirming that, although it is weaker than in the single-task condition, the GPE is present under divided attention. However, the interaction between serial position bin and trial type was not significant, *F*(2, 16,771) = 0.13, *p* = .880, η²_p_ < .001. This result suggests that the GPE is similar across cue conditions.

#### Alpha Power Across Task Demands

We hypothesized that alpha power would rise over the course of the list as memory load increased, based on prior findings (Sederberg et al., 2006). Using ANOVA on an alpha-specific model, we found a significant main effect of serial position bin *F*(2, 25935) = 151.53, *p* < .001, η²_p_ = .01. Post-hoc contrast analysis confirmed significant increases in alpha across the list for most serial position bin pairs across both task conditions, all z’s <-4.332, p’s <.001. However, there was no significant difference between the final two bins (middle-late bin vs. late bin, z=0.26, p=.796), suggesting that alpha power plateaus towards the end of the list.

There was a significant interaction between serial position and task *F*(2,25935)=4.48, *p*=.011, η²_p_ = .0003. However, the linear-trend analysis across serial-position bins indicated that the estimated slope of alpha power did not differ between task conditions, Single − Dual slopes β < −0.001 (SE = 0.003), *z* = −0.20, *p* = 0.84. Instead, direct bin-wise contrasts revealed significant task-related differences in alpha power at the early-middle bin, *z* = 2.46, *p* = .01, and late bin, *z* = –2.06, *p* = .04. The interaction thus appears to indicate a later increase and higher plateau in alpha power under dual-task rather than single task conditions.

#### Alpha Power By Cue Condition

In a second analysis restricted to dual-task trials, we tested whether changes in alpha power across serial position differed across cue conditions. However, the results indicated that the effect of target detection on alpha power was comparable across serial position bins, *F*(2,16679) = 127.32, *p*<.001, η²_p_ = .01. These results are consistent with additive effects of target detection and cognitive load.

## Discussion

The current study extends current literature by examining changes in EEG alpha and gamma band activity over multiple timescales. We examined how EEG activity responds to changing cognitive demands by having participants encode lists of words either under full attention or while concurrently performing a target detection task. Focusing on gamma and alpha bands, both known markers of attention processing, we examined how these frequencies changed as words were presented in a list. We further tested whether momentary attentional orienting to targets interacts with gradually increasing cognitive load as the list presentation unfolds.

We observed elevated high gamma band power at early list positions which declined later in the list, thereby replicating the gamma primacy effect (Sederberg et al., 2006; Serruya et al., 2014). This effect was attenuated under dual-task encoding, consistent with the idea that gamma reflects a sustained and focused encoding state that weakens as individuals divide their attention across items or tasks (Broitman et al., 2025; Broitman & Swallow, 2025; Sederberg et al., 2006).

As reported in previous studies, alpha activity increased across early serial positions (Healey & Kahana, 2020; Sederberg et al., 2006; Serruya et al., 2014), a pattern thought to reflect growing working memory load as cortical networks inhibit the processing of incoming stimuli and prioritize internal representations (Jensen & Mazaheri, 2010; Klimesch et al., 2007). Under divided attention, however, this rise in alpha was smaller early in the list, indicating that multitasking slows changes in both alpha and gamma activity during word list learning.

In a novel finding, alpha power was consistently lower with target detection despite simultaneously and additively increasing across serial position bins. This suggests that alpha power captures task-related processes at multiple timescales. Specifically, alpha power may capture a variety of cognitive demands, increasing gradually as the ability to encode new words decreases across a list presentation, yet rapidly increasing or decreasing in response to momentary task demands. This is consistent with recent work showing that target detection enhances memory uniformly across list positions (Broitman & Swallow, 2024), and with the theory that orienting and the underlying noradrenergic responses to targets are highly temporally constrained (Aston-Jones & Cohen, 2005; Swallow et al., 2022). By contrast, gamma power was not modulated by targets, suggesting that gamma reflects sustained attentional engagement rather than more transient orienting effects.

The attenuated changes in gamma throughout list presentation under dual-task encoding could arise from several mechanisms. One possibility is that the secondary task prevented attentional waning toward the end of the list by promoting vigilance and continued engagement throughout the presentation period, keeping spectral gamma power high. If so, then divided attention could paradoxically impair memory while stabilizing task engagement over time. This theory is supported by the relative stability in recall performance across serial position bins we observed among dual-task participants, compared to single-task participants. Alternatively, spectral power may have been lower during dual-task encoding at the onset of the list, reflecting greater initial levels of cognitive load and leading to less of a reduction over serial positions in the dual-task. Future studies should aim to manipulate task demands within subjects to adjudicate between these possibilities. It will also be important to examine whether differences in aperiodic EEG components contribute to the observed effects, given the growing interest in the influence of 1/f dynamics on spectral activity (Donoghue et al., 2020).

Some features of the current study’s design constrained our ability to compare the dynamics of neural activity across task demands. First, list lengths differed across task conditions, which balanced memory performance but complicates the direct comparison of serial position effects. We mitigated this by analyzing quartiles across conditions, but future studies should implement equal list lengths. Second, the between-subjects manipulation of task demands (dual vs. single), while necessary to avoid contamination from target detection instructions, limited sensitivity to subject-level differences. A within-subjects design that addresses possible carry-over effects should be developed to provide clearer contrasts. Finally, because target detection required a button press, some of the spectral power differences may reflect motor activity. Future studies may address this possibility by implementing non-motor secondary tasks (e.g., silent counting) to isolate cognitive factors.

## Conclusions

Our findings illustrate how transient and sustained changes in task demands impact neural correlates of attention during word list learning. Gamma power decreased over serial positions but this change was less steep under divided attention, potentially reflecting disrupted early engagement and prolonged task vigilance. Alpha power exhibited both gradual changes across list positions and rapid decreases following target presentations, reflecting its dual role in reflecting phasic and sustained attention processes. Together, these results highlight the differential roles of gamma and alpha in capturing multiple aspects of attention during memory encoding.

## Acknowledgements

The authors would like to thank Amy Krosch for allowing us to use her EEG facilities and Sanweda Maghabin and Sahib Kaila for assistance with data collection on this project. Funding for this project was provided by the Cornell University College of Arts & Sciences.

## Notes

### Competing Interest Statement

The authors have declared no competing interest.

